# Equipment-Free Personal Protective Equipment (PPE) Fabrication from Bacterial Cellulose-Derived Biomaterials via Waste-to-Wealth Conversion

**DOI:** 10.1101/2022.11.02.514716

**Authors:** Ramya Veerubhotla, Aditya Bandopadhyay, Suman Chakraborty

## Abstract

The recent COVID-19 crisis necessitated the universal use of Personal Protection Equipment (PPE) kits, generating tons of plastic wastes that inevitably lead to environmental damage. Circumventing the challenges stemming from such undesirable non-degradability on disposal, here we present an eco-friendly, robust, yet inexpensive and equipment-free method of growing biodegradable PPE fabrics by the fermentation of locally-sourced organic feed stocks in a rural livelihood. Using a pre-acclimatized symbiotic culture, we report the production of a high yield (up to 3.2 g fabric/g substrate) of bacterial cellulose, a biopolymer matrix, obtained by bacterial weaving. This membrane has an intricate, self-assembled, nano-porous 3D architecture formed by randomly oriented cellulose fibres. Scanning electron microscopy reveals that the pore size of the membrane turns out to be in the tune of 140 nanometers on the average, indicating that it can filter out viruses effectively. In-vitro results demonstrate assured breathability through the membrane for a filter thickness of approximately 5 microns. When subjected to soil degradation, the fabrics are seen to disintegrate rapidly and fully decompose within 15 days. With a favourable cost proposition of less than 1 US$ per meter square of the developed fabric unit, our approach stands out in providing a unique sustainable, and production-ready alternative to synthetic PPE fabrics, solving community healthcare and environmental crisis, and opening up new avenues sustainable under-served livelihood at the same time.

**Graphical abstract:** 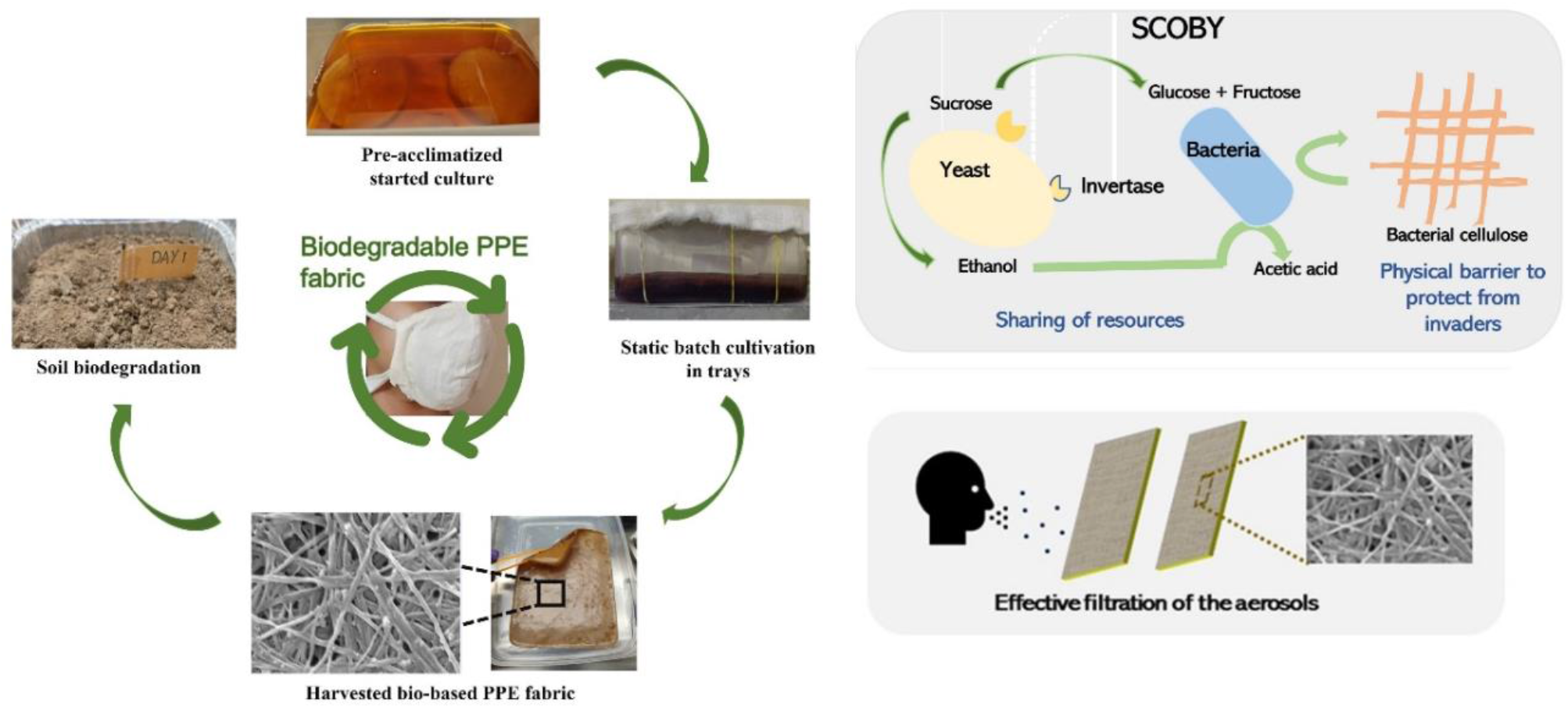

## 1. Introduction

Personal Protection Equipment (PPE) plays a pivotal role as a precautionary measure to arrest the spread of threatening infections transmitted by air-borne particulates. PPE primarily comprises face masks, coveralls, protective shoe cover, gloves, head caps, and several such items alike (Babaahmadi et al., 2021). They are expected to effectively safeguard the user from droplets or aerosols of the target pathogens (Ehsanifar et al., 2021). While the PPEs had remained to be on demand for long, they emerged to be effective life-saving items in a consumer supply-chain after the upsurge of the recent SARS-CoV-2 outbreak in different waves that created a serious shortage of their supplies globally, forcing several health workers to re-use otherwise single-use PPE components (Rowan et al., 2021). The World Health Organization (WHO) urged the manufacturing industries world-wide to escalate PPE production by at least 40 percent to cope with the critical PPE shortage (Singh et al., 2020). While the pandemic has been traversing its course to a naturally subsided endemic stage with significantly reduced threat, the need of such PPE kits tends to rise because of inevitable challenges from several different known sources of air-borne infections, as well as the possible emergence of newer variants of pathogens that may potentially disrupt human lives time and again (Facciola et al., 2021; Siwal et al., 2021).

Majority of the PPEs have ‘use-and-throw’ based disposability, generating massive amounts of unmanageable medical and environmental waste in the process (Adelodun et al., 2021). Considering Asia as a representative populated continent, around 2.2 billion face masks are being discarded every day after use. This contributes to thousands of tons of plastic waste ending up in the landfills (Allison et al., 2022). A large portion of such single-use plastics (mostly polypropylene based) interferes with water bodies and results in irreparable crisis to the aquatic ecosystems. This problem has been compounded by many folds due to the emergence of several other synthetic polymers such as polyamide, polyacrylonitrile, polyester, polycarbonate, nylon, polyethylene, polystyrene, etc., as commercially available alternate PPE fabric materials, benefitted by their low-cost, easy processing and low melt temperatures (Teymourian et al., 2021).The resulting non-biodegradability crisis, however, has risen to unbounded proportions as a consequence, fundamentally attributed to the disintegration of the polymeric fibres into microplastics (Shirvanimoghaddam et al., 2022; Masud et al., 2022; Noorimotlagh et al.,2021). As a specific example of the resulting environmental damage, recent studies indicated that untreated surgical masks made of polypropylene, when disposed on the topsoil, can take up to 28 years to fully biodegrade and result in long-term implications on human health that may induce livelihood adversities across the generations (Knicker et al., 2022).

The emerging trend of using polymers from alternative natural or semi-natural sources such as polyvinyl alcohol (PVA), polylactic acid (PLA), chitosan, chitin, gluten, alginate, cellulose acetate, and their composites for producing compostable PPE components has ushered immense promises of addressing the environmental challenge of producing completely biodegradable PPEs that may not lead to any environmental threat (Pandit et al., 2021; Chowdhury et al., 2020). These biopolymers may be used as the base materials to fabricate porous membranes/matrices that can effectively block the atmospheric contaminants. In a typical industrial set-up, this is accomplished by two common manufacturing technologies, namely, the spun-bound process and the melt-blown process. More recently, electrospinning and 3D printing technologies have advanced to an extent to provide viable rapid prototyping alternatives for generating micro/nanofibers of controlled dimensional features (Mishra et al., 2019; Swennen et al., 2020; De Sio et al., 2021; Gierthmuehlen et al., 2020). Some pertinent examples include: the use electrospinning technique to produce nanofibers made of bio-based, carbonized gluten material in combination with polyvinyl alcohol (PVA), and composites of bio-mineralized sponge spicules and PLA (Das et al., 2020; Wu et al., 2020). Such nanofibers are proven to block aerosol nanoparticles, favouring their use for face masks (Tebyetekerwa et al., 2020). More recently, the Fused Deposition Modelling technique has also been proposed as a promising technology for fabricating PLA-based 3D print protective masks (Vankova et al., 2020). However, these reported technologies of fabricating nano-fibrous PPE materials continue to get bottlenecked because of their extreme resource-intensiveness such as the demand of high electrical power and/ or complex setup, sophisticated instruments, and skilled human resources for the fabrication (Angammana et al., 2018).

Here, we report a pipelined material-to-product integration that deploys environmentally-friendly biopolymers on one hand and equipment-free fabrication paradigm on the other, simply by deploying activated microbes for both synthesizing and self-assembling the substrate into a porous nano-fibrous matrix, using their innate metabolic machines. The produced biopolymer matrix is shown to be directly usable as PPE fabric to sew facemasks, via architecting a desired nanofiber network with high precision by optimizing the microbial fermentation conditions. Proceeding further forward, we demonstrate the use of locally available, renewable organic feedstocks as substrates, to promote ‘value from waste’ and generate sustainable rural livelihood as well as good health at the same time.

## 2. Materials and methods

### 2.1 Pre-acclimatization of the starter culture

We use Symbiotic Culture of Bacteria and Yeasts (SCOBY), commercially purchased from Zoh Probiotics^®^, India, as the source of inoculum to perform microbial fermentation (Fig. 1). The mother culture is pre-acclimatized to pH value of 1 for growing the microbes selectively on tolerating highly acidic environments. The culture is maintained by repeated sub-culturing in infused tea solution, obtained by heating 6 g of black tea (Tata tea premium^®^), 50 g of sucrose (purchased from local market), and 0.4 mL of glacial acetic acid per litre.

**Fig. 1.**
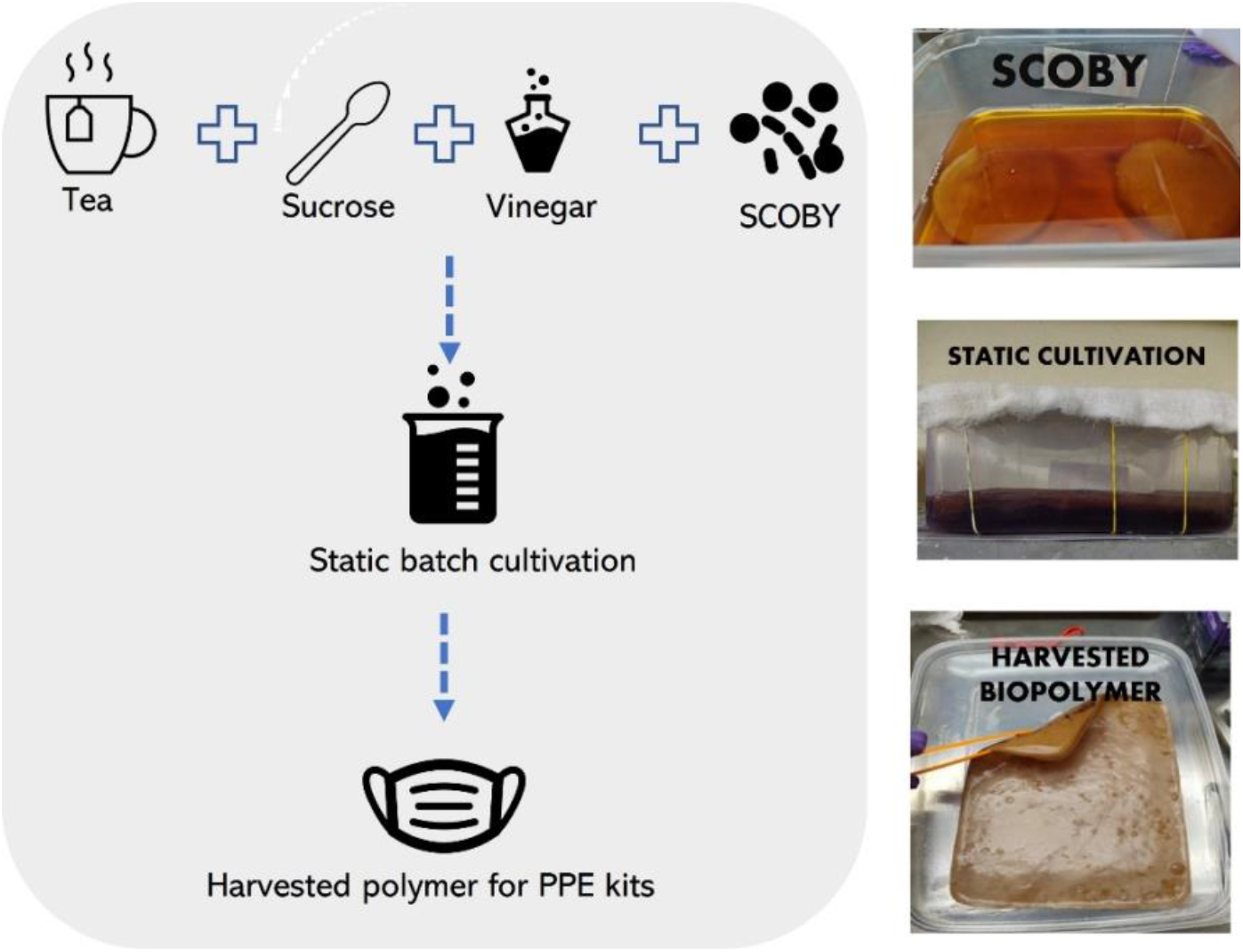
Schematic showing the steps involved in the biopolymer production process.

### 2.2 Static cultivation of the biopolymer and harvesting

Used black tea waste and sucrose are used as nitrogen and carbon sources in the production media respectively, 6 g of used black tea (per litre) and sucrose (25 g/L to 125 g/L) are brewed for 10 min to create a concoction. An initial pH of 1 is maintained by the addition of glacial acetic acid to the media. The cooled brew is then transferred to plastic food-grade rectangular trays for performing static fermentation. Each batch is inoculated with 30 mL of mother liquor and 25 cm^2^ of the microbial mat under sterile conditions. The trays are covered with multiple layers of sterile muslin cloth to allow oxygen penetration. The biopolymer floats at the air-liquid interface and is harvested after each batch run, as shown in Fig. 1. The harvested membrane is soaked in liquid soap solution for 48 hours to wash off impurities and cell debris. It is further coated with beeswax (0.01 g wax deposited per cm^-2^ of membrane) to modify the surface wettability. The resulting mats are then sterilized using UV radiation and stored for further use. For optimization studies, cell culture flasks (Thermo Scientific™ Nunc™ EasYFlask™ 75 cm^2^) are used.

### 2.3 Surface characterization by Scanning Electron Microscopy

Harvested samples are washed thoroughly with ethanol (sequentially increasing concentrations from 40 % v/v to 100 % v/v) and then fixed in 1.5 % glutaraldehyde solution for 1 h. Imaging is done to examine the surface morphology using Field Emission Scanning Electron Microscope (FE-SEM) [Zeiss Supra 40 FE-SEM] at 5 keV.

### 2.4 Biodegradation assessment

The used bio-fabric developed by the above process (3cm × 3 cm × 100 μm films) is biodegraded in known amounts of the topsoil at 35 °C for a period of two weeks. Samples are buried at a depth of 1 cm from the surface to allow the inherent soil microbial community for degrading the membrane under natural conditions. Throughout the experimental run, the soil is kept moist to facilitate degradation. At regular intervals, the samples are collected, washed thoroughly, and dried prior to the estimation of dry weight. Biodegradation is estimated as the percentage of weight loss of the degraded sample compared to its original weight as a function of time.

### 2.5 Pore-size analysis and flow rate calculation

From the SEM images, the pore sizes and their distributions are estimated by image analysis (using Matlab^®^ software). Considering each pore to be approximately cylindrical and subject to laminar, fully developed flow, the mean air flow velocity through each pore may be estimated as: 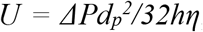, so that the volume flow rate in the direction of the pressure gradient, *ΔP*, reads: *Q = mAU*, where *dp* represents the average pore diameter, h represents the thickness of the mask/membrane, *η* (= 1.983 Ns/m^2^) represents the dynamic viscosity of air under the operating conditions, *m* is the fraction of pore area to the total area, *A* is the total area of the mask/membrane (=0.04 m^2^ for the experiments reported here), and *Q*, corresponds to the average volumetric flow rate. Thus, one gets: 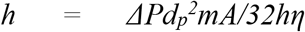, where *h* corresponds to the thickness of the membrane (Chowdhury et al., 2021).

## 3. Results and discussion

### 3.1 Process inventiveness

Industrial production of any bio-based product involves several upstream and downstream processing steps. Among them, raw materials, energy-intensive sterilization procedures, and complex product purification methods are the primary cost drivers (Wheelwright et al., 2020). In the case of bacterial cellulose (BC) production, 60 % of the total expenditure is solely contributed by the media components (Hussain et al., 2019). To render the present BC fermentation process not merely scientifically viable but also techno-economically competitive, the following inventive approaches are adopted.

#### 3.1.1 Locally available organic wastes as substrates for growing the biopolymer

Unlike the standard Hestrin–Schramm (HS) medium that consists of expensive nitrogen sources such as peptone and yeast extract, we use black tea waste to support the growth of microbes (Castro et al., 2012). Used tea waste, which is a common degradable out of domestic and beverage industry waste, in combination with sucrose, is shown to yield a higher amount of BC as compared to other organic wastes tested in the study (fruit peels, beverage wastes and inorganic nitrogen salts; data not shown). This is because, tea is rich in exclusive minerals (Cu, Fe, Mn, Ni and Zn), anions (fluoride, chloride, nitrates and sulphates), polyphenols, acetic acid, vitamin C, B, and caffeine (Yim et al., 2017). These compounds positively stimulate cellulose production by preventing c-di-GMP from being destroyed by phosphodiesterases (Jayabalan et al., 2010; Jayabalan et al., 2014). Tea polyphenols also exhibit anti-bacterial, antioxidant and antiviral activity (Mhatre et al., 2021). Therefore, it selectively inhibits the growth of several contaminants. It also lowers the pH, thus providing favourable fermentation conditions for the increased growth of acetic acid bacteria (AAB). However, our controlled experiments reveal tea concentration > 6 g/L inhibiting AAB adversely.

#### 3.1.2 Robust, stable, mixed culture as inoculum, pre-acclimatized to survive in hostile environments

Several pure strains of AAB such as *Acetobacter xylinum*, *A. xylinoides*, *Bacterium gluconicum*, *A. aceti*, *A. pasteurianus* and *Komagataeibacter* sp. are known to produce high amounts of BC (Dufresne et al., 2000). However, supporting the growth of such pure cultures requires controlled maintenance of sterile conditions throughout the production process. Herein, a selective, pre-acclimatized microbial culture, is used. Mixed cultures are highly robust; they facilitate better utilization of complex substrates and are highly resistant to environmental fluctuations. In the present approach, this provides a decisive advantage, since the multi-species consortia used contain both AAB and yeasts co-existing in strong symbiosis. This efficient division of labour among the yeast cells and AAB in Symbiotic culture of bacteria and yeast (SCOBY) enable better utilization of the nutrients as well as provide enhanced protection from invading agents, resulting in favourable resource storage in the form of BC (May et al., 2019).

Fig 2 (a) schematic represents strong symbiosis in SCOBY and AAB on the surface of BC material in the inoculum. A detailed description of the symbiotic interaction in the mixed culture is outlined in Section S1 of the Supplementary Information (SI). Strategic enrichment of the consortia at a very low pH ensures enrichment of BC-producing AAB stains that can tolerate hostile environments. Adopting low pH can further reduce the risk of contamination significantly, eliminating the need of maintaining stringent sterile conditions during the fermentation process. Fig. S1 of the SI shows the optimal production of biofabric by the consortia at different pH, following acclimatization.

**Fig. 2.**
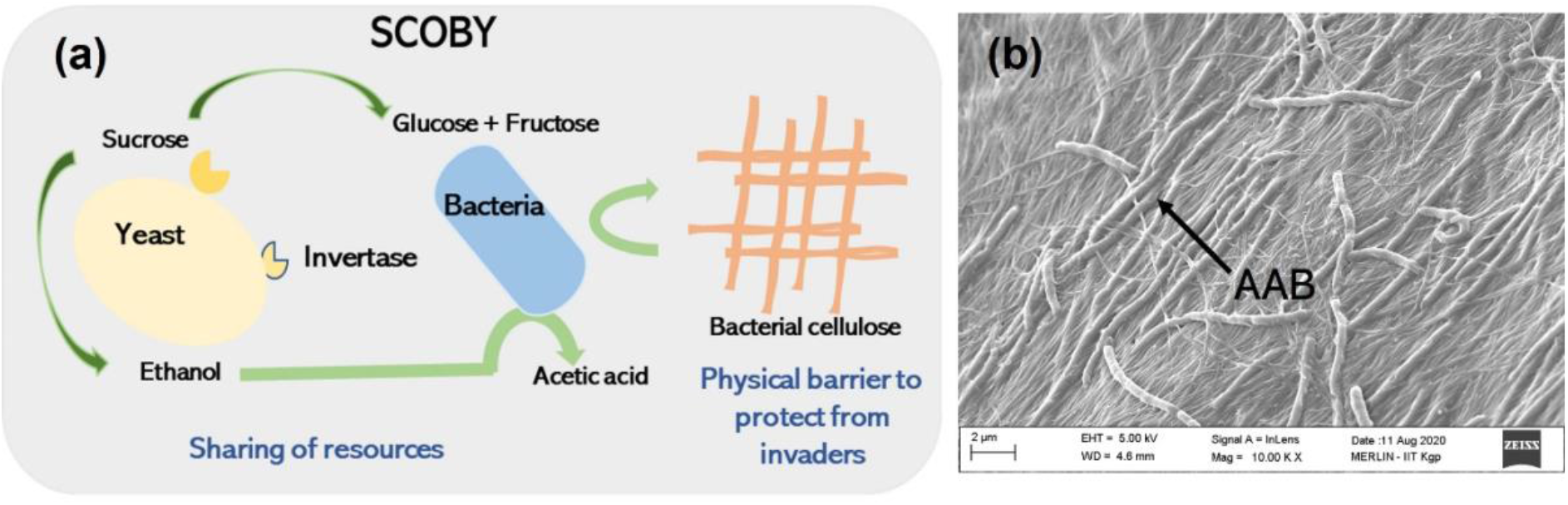
(a) Interaction of microbes in SCOBY and (b) SEM images of the BC mat showing colonization of AAB on the surface.

#### 3.1.3 Simple downstream processing

Bacterial cellulose is an extracellular polymer secreted by AAB into the media. As a result of yeast metabolism, carbon dioxide bubbles, accumulated in the media, cause the BC mat to float on the surface. Therefore, it can be collected easily. Further purification and disinfection are done by soaking in an eco-friendly soap solution and ultra-violet irradiation, respectively. Such hassle-free, simple downstream processing further reduces the overall expenditure of the process.

### 3.2 Selecting the material and the process parameters

Properties of BC such as pore size, shape, volume distribution, and optical transparency can be tuned by growth media components. BC is also known for its hydrophilicity and high tensile strength which can be tailored by both in-situ and ex-situ methods (Ashrafi et al., 2019; Tang et al., 2010). This renders BC one of the ideal materials for fabrication of PPE, as its properties can be customized as per the needs.

Optimization studies on the BC matrix are performed in set flaks by varying the sucrose concentrations from 25 g/L to 125 g/L at pH 1 (Fig. 3(a) and 3(b)). Results indicate that 75 g/L sucrose concentration comes up with an unprecedented maximum yield of 3.2 g biofabric/ g sucrose (corresponding to 23.4 g/L of culture media) in the batch mode; see Fig. 3(c). To the best of our knowledge, this is one of the highest yields reported for bacterial cellulose production (Goh et al., 2012; Cakar et al., 2014; Raiszadeh-Jahromi et al., 2020; Singhsa et al., 2018; Ruka et al., 2012; Shezad et al., 2009). Such high yields can be attributed to potent BC-producing microbial populations in the mixed culture used. To arrive at the maximum yield, the batch fermentation is run for 50 days to ensure complete substrate utilization. However, the profile of the biopolymer assimilation indicates that most of the polymer production occurs within a period of 21 days.

**Fig. 3.**
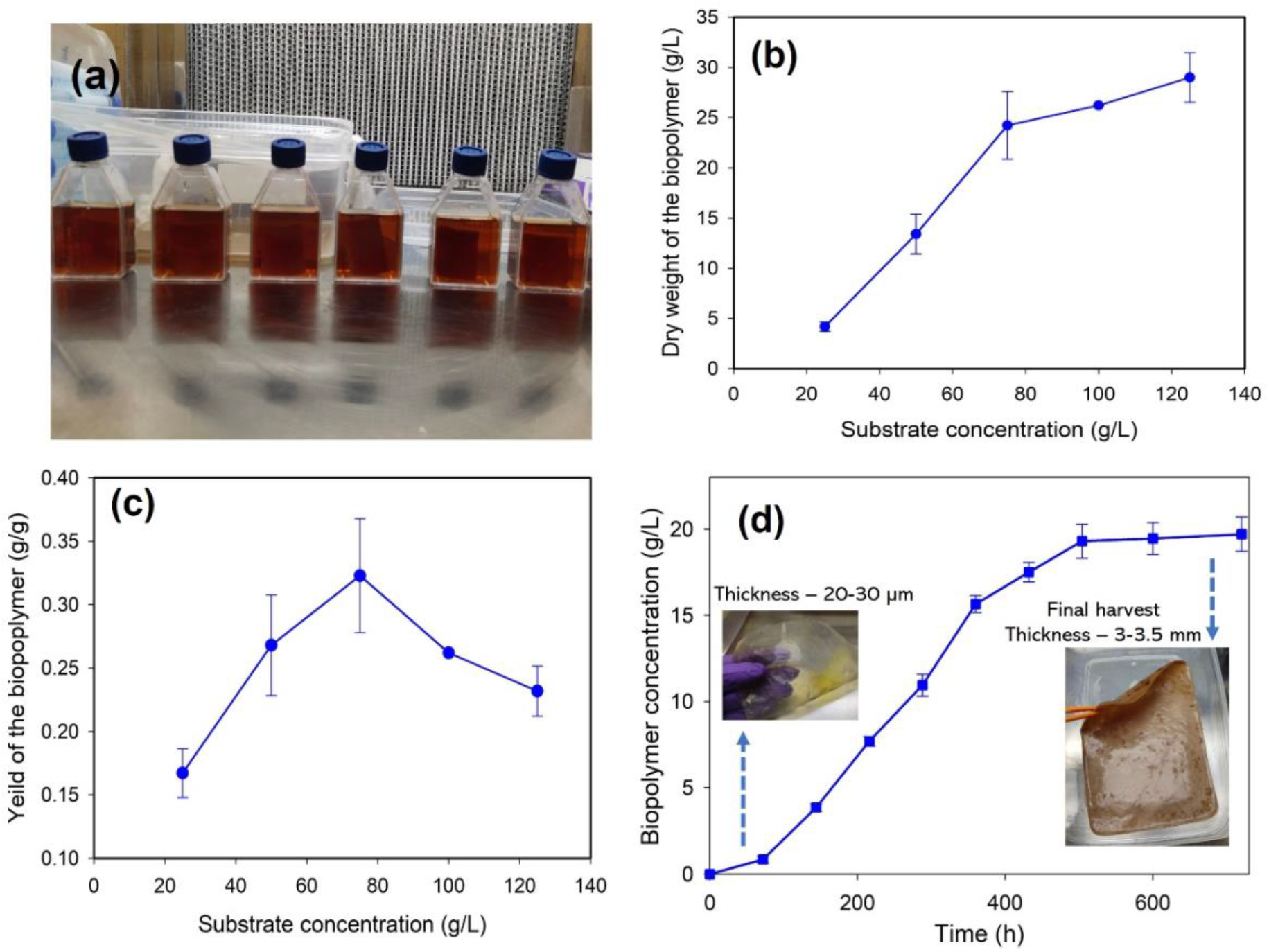
(a) Set flasks inoculated with mixed microbial culture to perform the optimization studies (b) Amount of the biofabric obtained as a function of the substrate concentration (c) yield versus sucrose concentration and (d) profile of polymer assimilation with time.

The BC mats harvested at end of day 2 are optically transparent having a thickness of 20-30 μm; see Fig. 3(d). This can be an added advantage in developing transparent masks, benefiting deaf-mute people who rely on facial expressions for communication (Avossa et al., 2021). Thickness of the mats also decides the air flow rate across mats and is strongly dependent on substrate concentration as well as the oxygen diffusion. By optimizing the feeding strategy and the substrate concentration, one can get the membranes of desired thickness in the same batch. Fig. S2 of the SI shows the thickness of mats as a function of substrate concentration. The biofabric production is found to yield a maximum of 85.4 g in 100 days (corresponding to a wet weight of approximately 650 g) when intermittently fed with media every 20 days. Fig. S3 of the SI shows the production of the BC in fed-batch mode of operation. The biopolymer pellicle formation in different layers is an indicative of the feeding cycle, as shown in Fig S4 of the SI.

### 3.3 Effectiveness against the virus particles

FE-SEM imaging (Fig. 4a) reveals randomly oriented 3D nano-fibrous architecture of the cellulose matrix in the biopolymer that is intrinsically woven by the microbes. The average pore size of the BC membrane as estimated from the SEM images turns out to be around 140 nm. This is in the tune of the diameters of typical virions (Bar-On et al., 2020). This implies a favourable proposition of blocking the virus particles by simply offering a steric hindrance against their suction into the respiratory pathways via human nostril. Typically, in case of BC, the substrate chosen, growth conditions, and post-harvesting strategy decide the pore size and thickness of the resulting membrane. Reported studies indicate that BC can be produced in different average pore sizes ranging from 1 μm down to 4 nm (Gao et al., 2011). Thus, ultra-narrow porous pathways can be fabricated on the same substrate material, resulting in tunable resistance against foreign particle penetration as per the particular particle size.

**Fig. 4.**
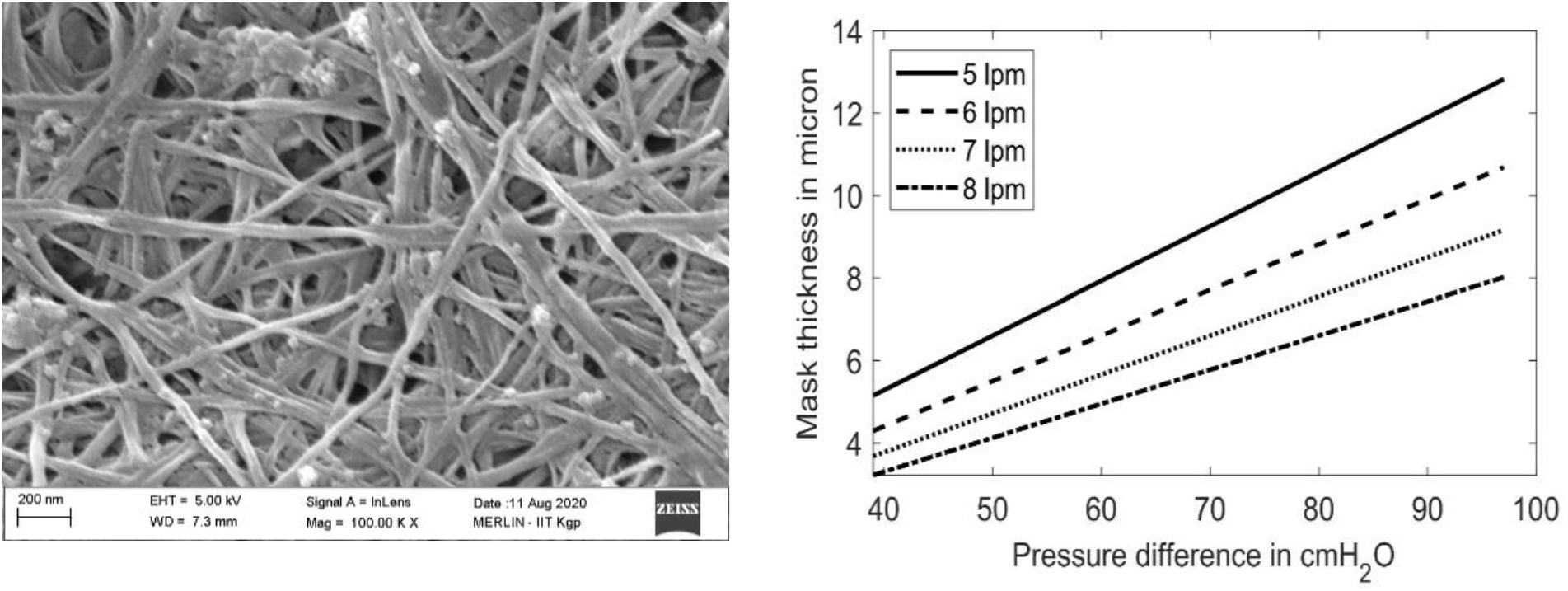
(a) Scanning electron microscopy (SEM) of the treated biopolymer filter at a magnification of 1,00,000 X and (b) Membrane thickness as a function of the pressure difference for different air flow rates.

### 3.4 Breathability

Fig. 4(b) represents a plot of the membrane thickness as a function of ΔP for different flow rates, following the design criterion as outlined in Section 2.5. Lausted et al. reported that, among women, the inspiratory pressure can vary from 68 cm H_2_O to 28 cm H_2_O, whereas the maximum expiratory pressure can vary from 39 cm H_2_O to 61 cm H_2_O (Lausted et al., 2006). For men, these parameters vary from 39 cm H_2_O to 97 cm H_2_O and 63 cm H_2_O to 97 cm H_2_O, respectively. Further, whereas the peak flow rate may range up to 600 litre per minute (lpm) for fit young individuals, the same under normal conditions may average around 8 lpm (Dikshit et al., 2005). Thus, considering an average ΔP of 60 cm H_2_O and flow rate of 8 lpm, we may conclude that a mask of 5 μm thickness can sufficiently ensure easy breathability.

### 3.5 Biodegradation

When subjected to soil degradation under controlled conditions, it is observed that the material significantly loses its weight within 12 days. Complete biodegradation in the soil occurs within two weeks as shown in Fig 5. Since soil is a good source of cellulose degrading microbes, no additional bacterial cultures or enzymes are added to degrade the fabrics. As compared to other biopolymer membranes such as polyhydroxy butyrate and polylactide, the BC films have a faster degradation rate (Tokiwa et al., 2006; Altaee et al., 2016). The degradation can be further promoted by the addition of soil microbes such as *Bacillus* sp. and *Rhizopus* sp. (Zahan et al., 2020). Thus, the filters developed here can be safely and rapidly composted.

**Fig. 5.**
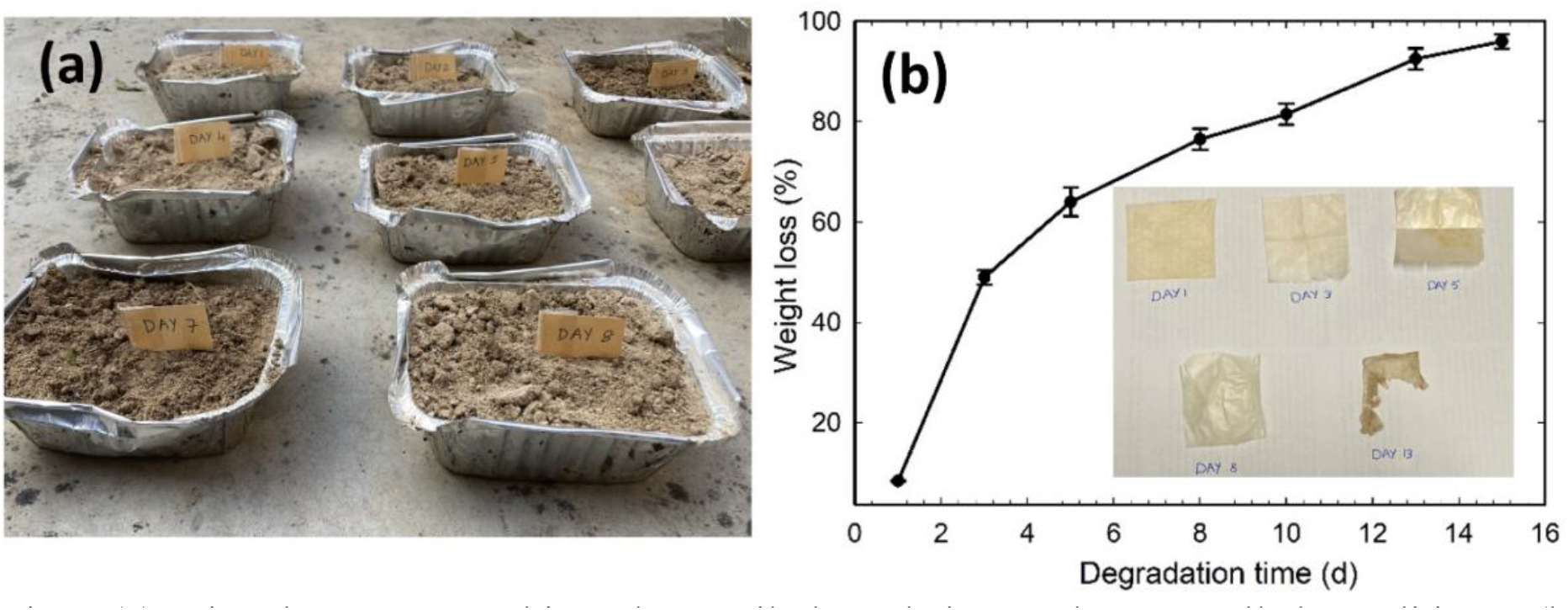
(a) Biopolymer mat subjected to soil degradation under controlled conditions, (b) degradation profile. Inset shows mats recovered from soil after 1, 3, 5, 8 and 13 days, respectively.

## 4 Economic value proposition and prospect for use

Table 1 provides an estimate of the total expenses incurred for bioconversion of organic wastes to BC. Our analysis indicates that the 1 m^2^ of the biofabric (of about 100 grams per square meter) can be produced at a cost as low as 1 $, requiring approximately 25 L of production media and inoculum volume of 1.25 L to set-up the fermentation. Use of a mixed culture, harsh fermentation conditions, and renewable raw materials are responsible for such dramatic cost reduction as compared to the state of the art. Thus, the use of BC for PPE manufacturing is both economically profitable and environmentally sustainable. Scale-up and further automation of the process can further minimize the costs, indicating promising commercialization prospects.

**Table 1.**
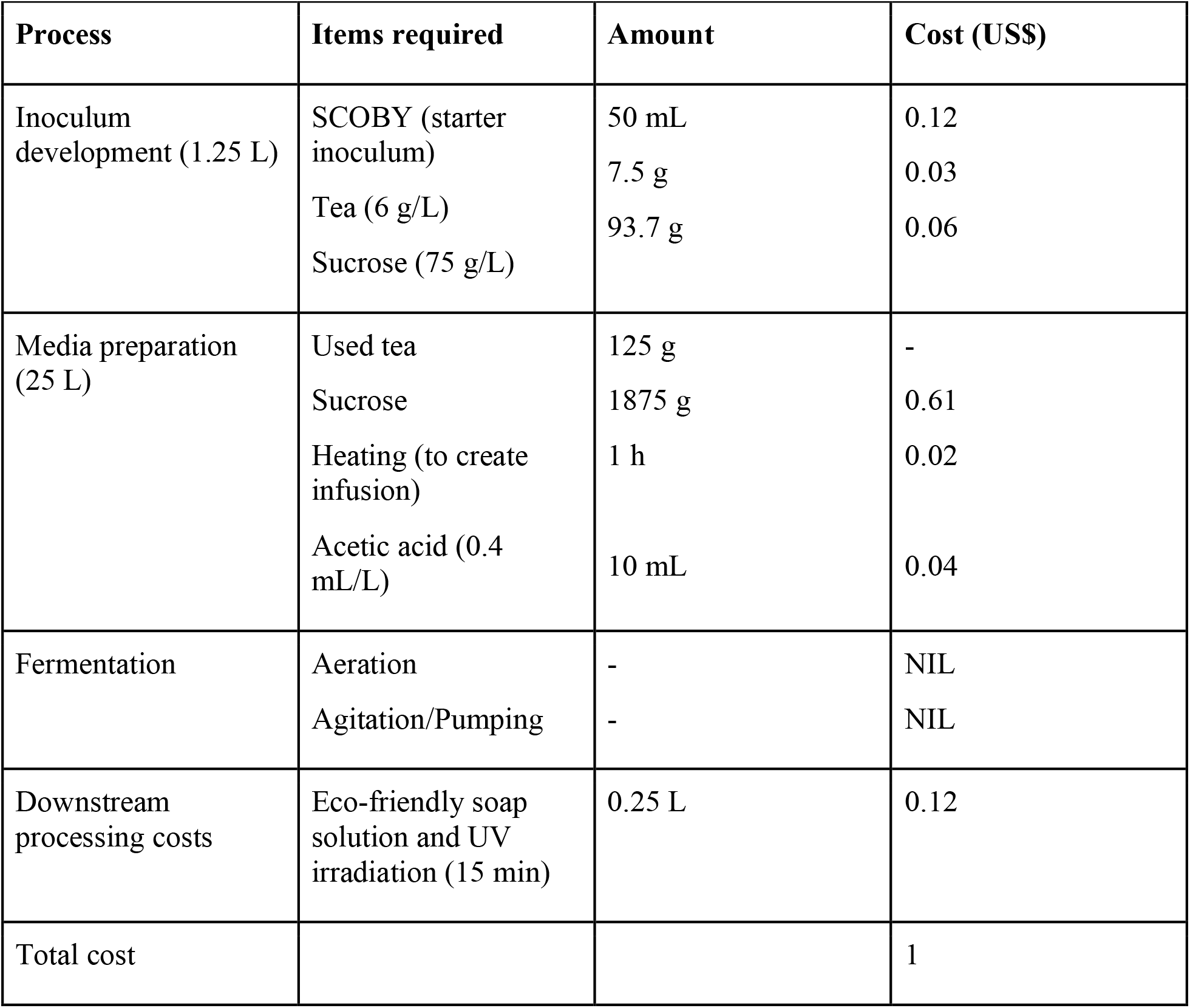
Cost analysis for producing 1 m^2^ of the biofabric produced using static fermentation

| Process | Items required | Amount | Cost (US\$) |
| --- | --- | --- | --- |
| Inoculum development (1.25 L) | SCOBY (starter inoculum) | 50 mL | 0.12 |
|  | Tea (6 g/L) | 7.5 g | 0.03 |
|  | Sucrose (75 g/L) | 93.7 g | 0.06 |
| Media preparation (25 L) | Used tea | 125 g | - |
|  | Sucrose | 1875 g | 0.61 |
|  | Heating (to create infusion) | 1 h | 0.02 |
|  | Acetic acid (0.4 mL/L) | 10 mL | 0.04 |
| Fermentation | Aeration | - | NIL |
|  | Agitation/Pumping | - | NIL |
| Downstream processing costs | Eco-friendly soap solution and UV irradiation (15 min) | 0.25 L | 0.12 |
| Total cost |  |  | 1 |

The material fabricated here can be sewn into different PPE components such as face masks, coveralls, and shoe covers. It can either be integrated into the existing fabric masks or N95 respirators to obtain high filtration efficiency (Prototype demonstrated in Fig. 6) or be used as stand-alone material instead of other medical fabrics. In-situ addition of certain anti-pathogen compounds to the biofabric can further improve its functionality by deactivating the pathogen immediately upon exposure (Wu et al., 2014; Hewawaduge et al., 2021).

**Fig. 6.**
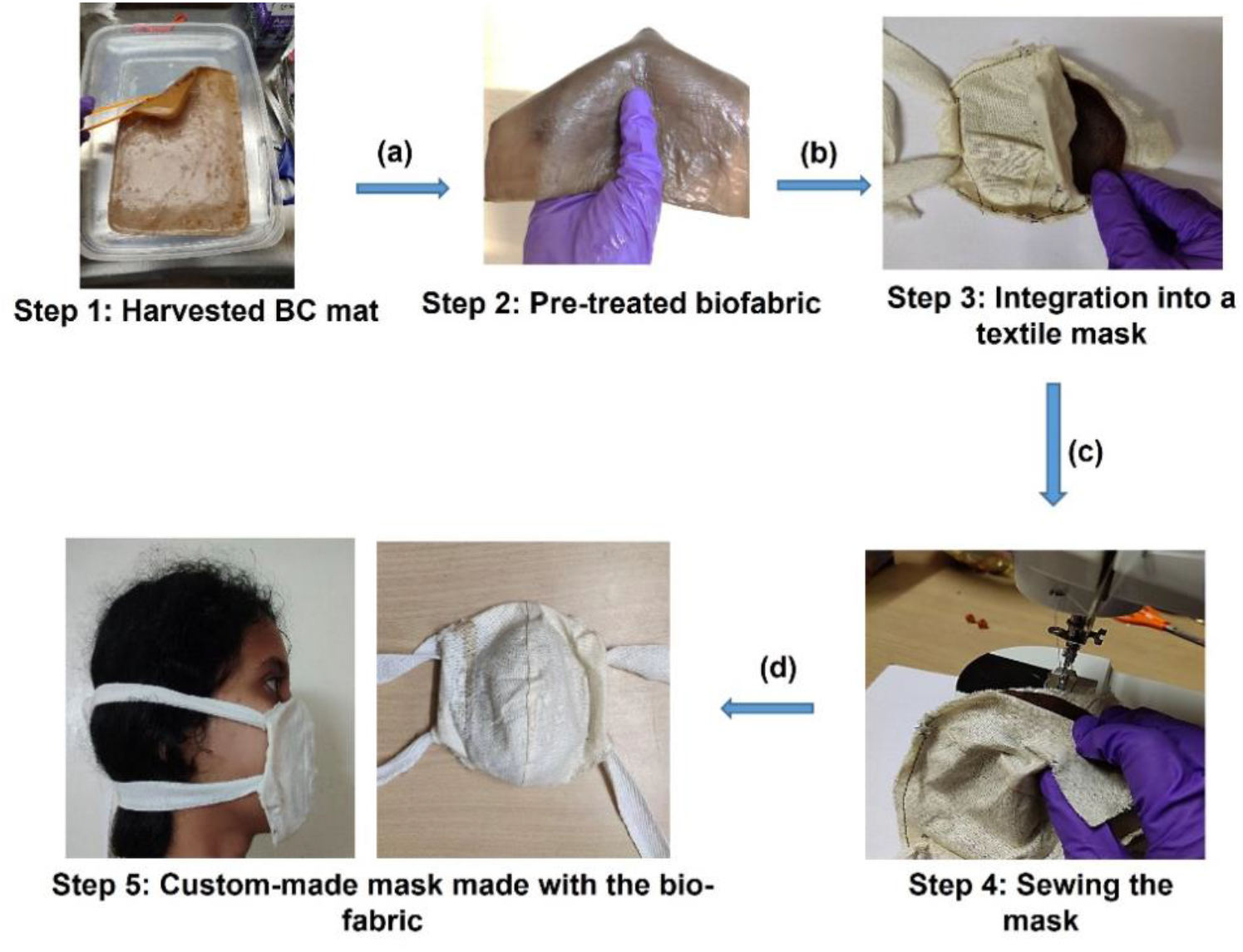
Prototype of the biodegradable PPE mask developed in this work.

## 5 Conclusions

We demonstrated a facile, cost-effective method to produce biodegradable PPE fabrics that relied on the naturally occurring metabolic processes of the microbes. Circumventing the resource-intensive routes of electrospinning and 3D printing for developing precision naoporous biomaterials, the present technology was demonstrated to be equipment-free and inherently economized. Such unique value propositions could be realized by the use of organic wastes, robust mixed microbial culture, and hostile fermentation conditions (low pH), even under unsterile conditions. The resulting process flow succeeded in dual prong reduction of the overall cost (~1 $/ m^2^), along with a concomitant high production yield of 3.2 g biofabric/ g sucrose. Further, the biopolymer-based material could readily be customized for fabricating coveralls, face masks, shoe covers, and several other PPE items. The resulting paradigm of the use of indigenous resources, process economization via inventive protocols involving waste material recovery and supreme eco-friendliness with no compromise in the performance efficacy as well as fabrication scalability appears to be ideal for personal protection against dynamically evolving infection threats via airborne particulates, ensuring sustainable development and good health and well-being for all at the same time.

## Abbreviations

PPE: personal protection equipment
BC: bacterial cellulose
SEM: scanning electron microscopy

## Author statement

RV did the experiments and writing. AB did the analysis and problem formulation. SC did the problem formulation and writing.

## Acknowledgments and Funding

SC acknowledges SERB, Department of Science and Technology, Government of India, for Sir J. C. Bose National Fellowship. RV and AB acknowledge funding and facilities of the MN Faruqui Innovation Center at IIT Kharagpur.

## Data availability

Data can be made available upon request.

