## Supplementary Information for "Equipment-Free Personal Protective Equipment (PPE) Fabrication from Bacterial Cellulose-Derived Biomaterials via Waste-to-Wealth Conversion"

**Ramya Veerubhotla[1], Aditya Babdopadyay[2], and Suman Chakraborty[2]**

**[1]Aarhus University Centre for Water Technology (WATEC), Department of Biology, Section for Microbiology, Aarhus University, 8000, Aarhus C, Denmark.**

**[2]Department of Mechanical Engineering, Indian Institute of Technology Kharagpur Kharagpur-721302, INDIA**

### S1. Mechanism of species-level interaction among yeasts and Acetic Acid Bacteria in SCOBY

Yeasts in the mother liquor break down sucrose to its monomeric forms with the help of invertase. Simultaneously, they convert a portion of these sugars to alcohol and carbon dioxide. The metabolic products of yeasts are taken up by AAB and used as a source of energy. AAB also assimilate the simple sugars and polymerize them through β-1→4 linkages, producing thick BC pellicles at the air-liquid interface. The BC mat acts as a strong barrier minimizing the penetration of invading pathogens into the culture medium, thereby protecting the underlying yeast cells from radiation, antibiotics, molds, and other contaminants. As a result, BC renders to be naturally anti-fungal and anti-bacterial. The BC mat floats due to the carbon dioxide (in the form of bubbles) generated by yeast, allowing the aerobic AAB to embed in the porous matrix at the surface to breathe (May et al., 2019).

### S2. Optimal conditions for production

High production yield (around 28 g/L) is obtained when 100 g/L is fed to the system in the second feeding cycle. Likewise, when 50 g/L is fed in the third round, only 16 g/L of the polymer is produced, indicating that the amount of the polymer assimilated by the microbes is directly proportional to the substrate added. However, in the further feeding cycles, despite the addition of 75 g/L of sucrose to the culture media, the production drops subsequently, yielding only 9.7 g/L. This is probably due to non-availability of sufficient nutrients to the microbes in the mat due to poor diffusion. Thus, the fed batch mode of cultivation is only beneficial for at most three feeding cycles.

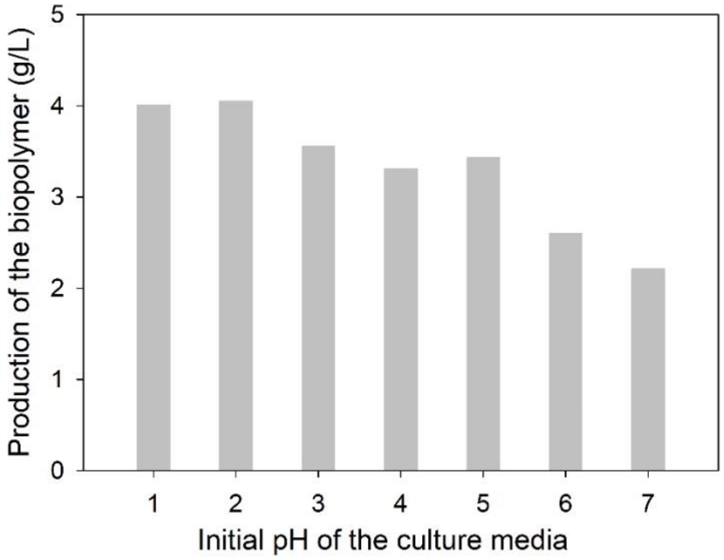


Fig. S1. Amount of the polymer produced by the pre-acclimatized mixed culture as a function of pH using tea infusion

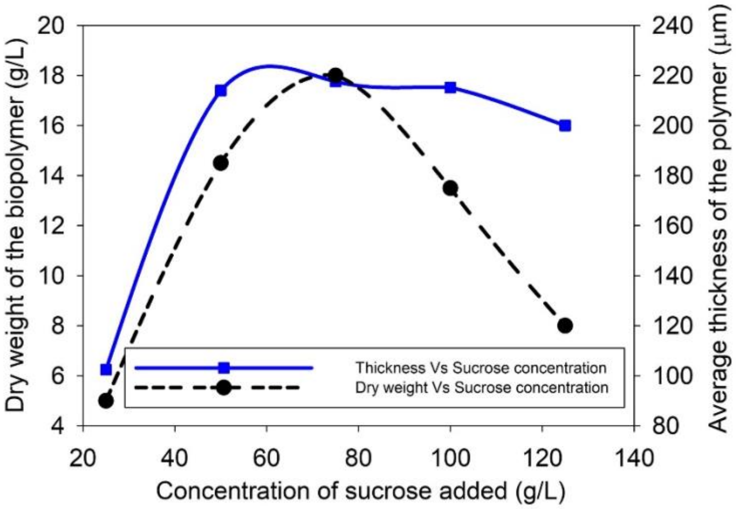


Fig. S2 Thickness of the BC mats as a function of substrate concentration (fermentation time of 21 days)

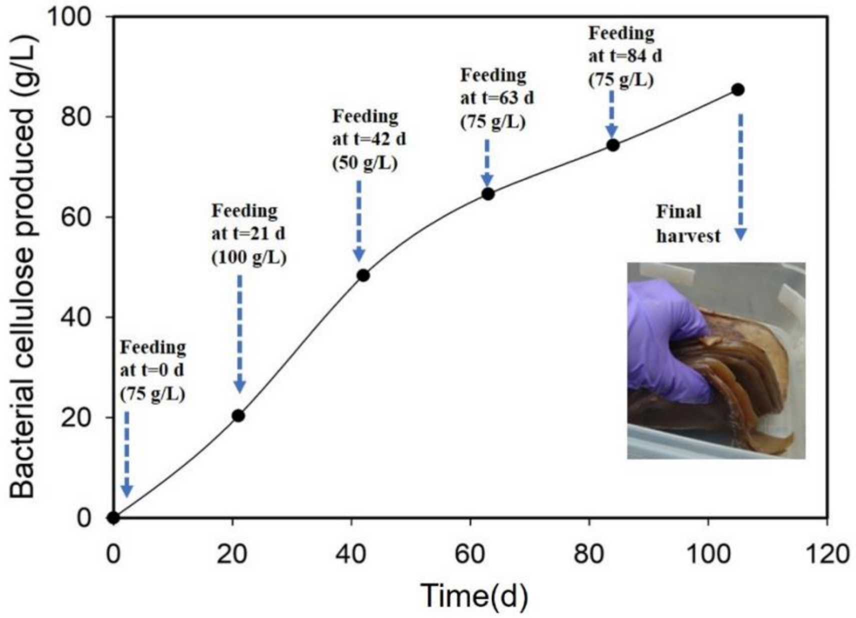


Fig. S3. Fed-batch cultivation of BC mats during a 100-day fermentation period

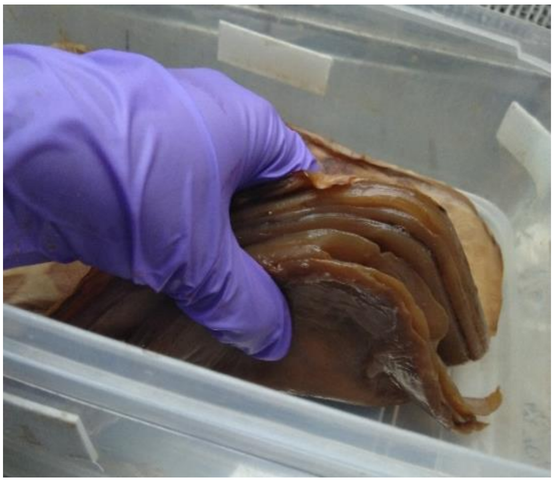

Fig. S4. Layer-by-layer assembly of BC, an indicative of the feeding cycle
